# Sensing of viral RNA in plants via a DICER-LIKE Ribonuclease

**DOI:** 10.1101/2023.01.10.523395

**Authors:** Carsten Poul Skou Nielsen, Lijuan Han, Laura Arribas-Hernández, Darya Karelina, Morten Petersen, Peter Brodersen

**Affiliations:** Copenhagen Plant Science Center, University of Copenhagen, Ole Maaløes Vej 5, DK-2200 Copenhagen N, Denmark; Max Planck Institute for Biology Tübingen, 72076 Tübingen, Germany; Department of Biology, University of Copenhagen, Ole Maaløes Vej 5, DK-2200 Copenhagen N, Denmark; Novo Nordisk A/S, Hagedornsvej 1, Gentofte, Denmark; Institut de Biologie Moléculaire des Plantes du CNRS, 12 rue du Général Zimmer, 67084 Strasbourg Cedex, France

## Abstract

Sensors of intracellular double-stranded RNA are central components of metazoan innate antiviral immunity, but such sensors have not been identified in plants. RNA interference (RNAi) constitutes a potent plant antiviral defense mechanism that relies on conversion of viral RNA into small interfering RNAs by two DICER-LIKE (DCL) ribonucleases, DCL4 and DCL2. Here, we show that while plant DCL4 is dedicated to RNAi, cytoplasmic dicing by DCL2 also triggers RNAi-independent defense gene expression via at least two intracellular nucleotide-binding domain/leucine-rich repeat (NLR) immune receptors. Combined *DCL4/NLR* inactivation abrogates basal resistance to a positive strand RNA virus. Our results redefine the basis of plant antiviral immunity, including autoimmunity as an explanation for DCL2-dependent growth arrest in *dcl* and RNA decay mutants in several plant species.

**One sentence summary:** The plant immune system uses Dicer-like ribonucleases for both antiviral RNA interference and double-stranded RNA sensing.

---

Recognition and elimination of potential pathogens of paramount importance for host survival and reproduction. Plants have an innate immune system with at least two distinguishable, but interconnected layers. In the first, conserved pathogen molecules activate receptor signaling to induce an immune state, referred to as pathogen-associated molecular pattern (PAMP)-triggered immunity (PTI) (*1*). Successful pathogens employ virulence factors, or effectors, to prevent or delay PTI, often by action inside host cells (*2*). In turn, a series of intracellular nucleotide-binding site-leucine-rich repeat (NLR) immune receptors encoded by *Resistance* (*R*) genes, constitute the core of a second layer of defense, effector-triggered immunity (ETI) (*1*). R proteins often recognize effectors indirectly through association with effector targets (*3*) and are activated by oligomerization to form resistosomes upon change in such host factors (*4, 5*). R proteins with N-terminal coiled-coil (CC) domains (CNLs) may insert into membranes and act directly as non-selective Ca^2+^-channels upon pentamerization (*6–10*), while R proteins containing N-terminal Toll/Interleukin-1 receptor (TIR) domains (TNLs) tetramerize (*11, 12*) to form an enzyme that cleaves NAD^+^ to produce a variant glycocyclic ADP-ribose with presumed second messenger activity (*13–16*). Mutation of effector targets may cause autoimmunity via constitutive R protein activation, a condition resulting in growth inhibition sometimes accompanied by programmed cell death and leaf chlorosis (*17–21*). Such mutations may also cause defective pathogen recognition if activation of the cognate R proteins requires a modified version of the host factor rather than its disappearance altogether (*22–25*).

In addition to proteins, nucleic acid components of viral intruders can be sensed. In mammals, cytoplasmic double-stranded RNA (dsRNA) induces antiviral immunity (*26*) via binding to the RNA helicase domains of Retinoic Acid Induced Gene-I (RIG-I)-like receptors (RLRs) (*27*). In plants, signaling from dsRNA sensors has not been described, as plants use RNA interference (RNAi) as a basal antiviral immune system which may be thought of as a PTI for viruses. In antiviral RNAi, viral dsRNA is processed into small interfering RNA (siRNA) by DICER-LIKE (DCL) ribonucleases that contain helicase domains closely related to those of mammalian RLRs. In turn, virus-derived siRNAs program endonucleolytic RNA Induced Silencing Complexes (RISCs) against viral RNA, and silencing is amplified by the RNA-dependent RNA Polymerase RDR6 recruited to RISC-targeted viral RNA, thus generating more viral dsRNA for processing via DCLs (*28*). In the current model of plant antiviral RNAi, two of four DCLs, DCL4 and DCL2, perform largely redundant functions in generation of antiviral siRNAs, and basal immunity against RNA viruses lacking anti-RNAi effectors is abrogated only when both *DCL4* and *DCL2* are missing (*29–31*). Nonetheless, DCL4 and DCL2 are biochemically distinct: DCL4 produces 21-nt siRNAs while DCL2 produces 22-nt siRNAs (*32, 33*). Crucially, most RNA virus-derived siRNAs are produced by DCL4 in wild type plants, (*29–31, 34*), and single mutation of *DCL4*, but not of *DCL2*, can partially break basal antiviral resistance (*31*). Moreover, in transgene silencing, DCL2 and DCL4 have distinct roles with particular relevance of DCL2 in RNAi amplified via RDR6 (*35, 36*). Most importantly, in the absence of viral infection, mutation of arabidopsis *DCL4* leads to growth inhibition, sometimes associated with leaf yellowing. This incompletely penetrant phenotype is fully suppressed by inactivation of *DCL2* (*36–38*), but is exacerbated by mutation of the enzyme required for microRNA (miRNA) biogenesis, DCL1 (*34*), or by mutants in cytoplasmic RNA decay (*39, 40*). Simultaneous inactivation of several cytoplasmic RNA decay factors is also sufficient to cause similar DCL2-dependent growth phenotypes (*39*). The growth inhibition is thought to result from generation of ectopic DCL2-dependent siRNA populations targeting mRNAs encoding the regulators of phloem formation and sugar metabolism, *SMXL4/5*, the nitrate reductase *NIR2* or the calvin cycle component trans-ketolase *TKL1* (*38, 39, 41, 42*). Although knockout of *SMXL4/5, NIR2* and *TKL1* does cause growth inhibition (*34, 38, 39*), proof of the causality of ectopic silencing of any of these factors for DCL2-dependent growth inhibition is missing.

## DCL-dependent activation of innate immunity is deeply conserved in plants

Since the reduced growth and leaf yellowing in *dcl4* mutants is reminiscent of autoimmune phenotypes (*43*), we tested whether the growth phenotype of *dcl4* mutants was associated with activation of canonical immune responses associated with ETI or PTI. Indeed, defense marker genes showed clear upregulation in *dcl4-15* mutants (Fig. S1), and expression profiles showed that gene ontology (GO) terms related to defense responses were enriched among genes upregulated in *dcl4-15* mutants exhibiting the growth phenotype (Fig. 1A). This defense gene expression profile was nearly fully erased upon silencing of *DCL2* with an artificial miRNA in *dcl4-15 (dcl4-15/amiR-DCL2*, Fig. 1B). Thus, loss of DCL4 in arabidopsis leads to activation of DCL2-dependent autoimmunity.

**Fig. 1.**
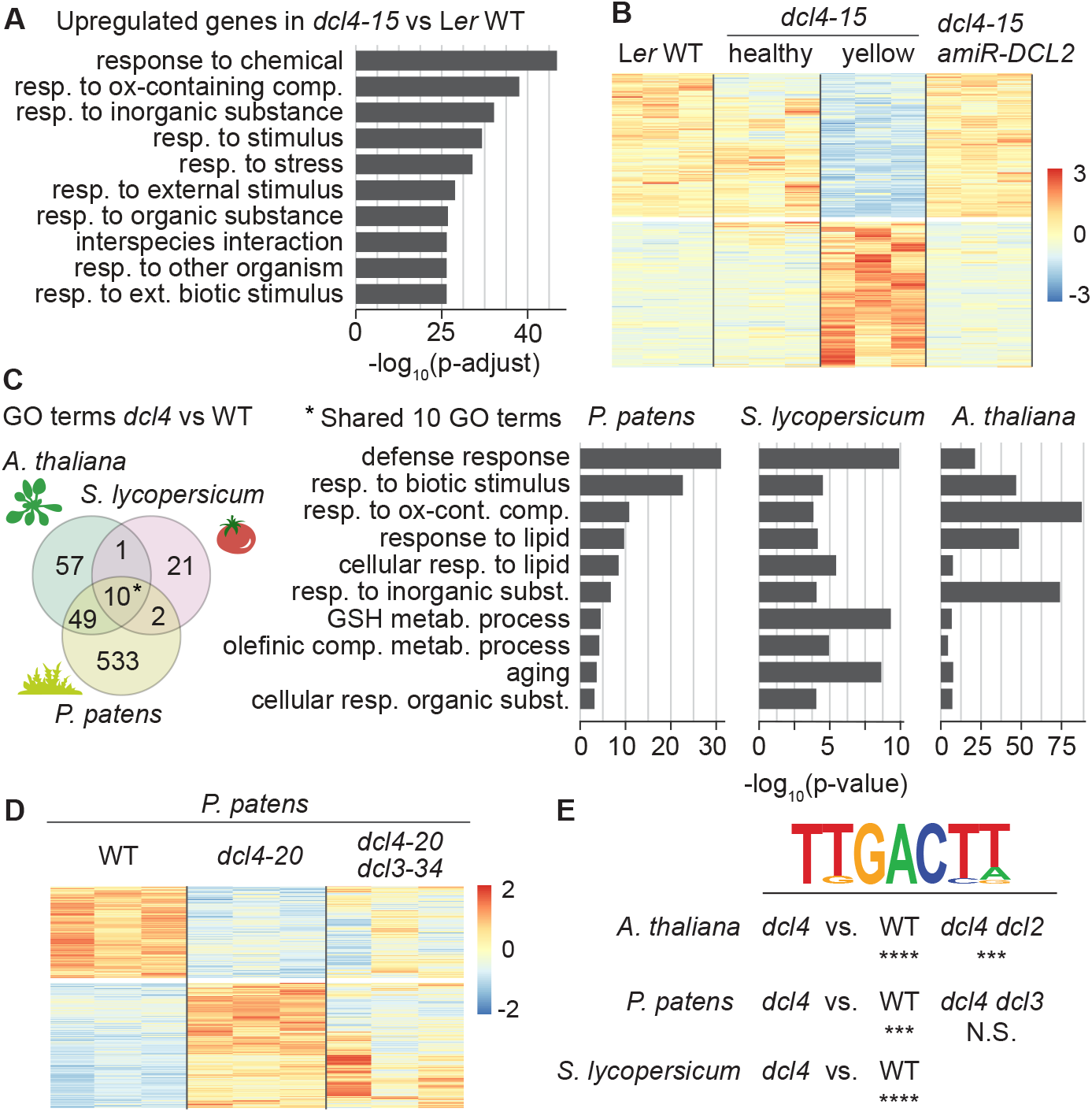
Knockout of *DCL4* causes autoimmunity in *Arabidopsis, Solanum lycopersicum* (tomato) and *Physcomitrium patens* (moss) (**A**) The 10 most significantly enriched gene ontology (GO) terms among genes upregulated in yellowing seedlings of *Arabidopsis Ler dcl4-15* compared to wild type, as determined by mRNA-seq analysis. Seedlings were harvested 21 days after germination. (**B**) Row-normalized heat map of differentially expressed genes as determined by mRNA-seq between *Arabidopsis Ler* seedlings of the following types: wild type (all green), *dcl4-15* (healthy and green), *dcl4-15* (yellowing), *dcl4-15 amiR-DCL2* (all green). The proportion of yellowing *dcl4-15* individuals was 50%, and was 0% upon *amiR-DCL2* expression, as reported in (). (**C**) (Left) Overlap between enriched GO terms among genes upregulated in *dcl4* mutants compared to wild type, in *Arabidopsis Ler* seedlings, *S. lycopersicum* leaves, and *P. patens* gametophores. (Right) List of the 10 GO terms that overlap among the three species (marked with an asterisk in the Venn diagram). (**D**) Row-normalized heat map showing differentially expressed genes in *P. patens* wild type, *dcl4-20* and *dcl4-20/dcl3-34*. (**E**) Enrichment of the W-box TTGACTT in promoters of genes upregulated in *dcl4* mutants compared to wild type in all three species, and to *dcl4/dcl3* for *P. patens* or *dcl4/dcl2* for *Arabidopsis*. Significance of enrichment as estimated by FIMO (see Methods), **** P < 0.0001, *** P < 0.001, NS, non-significant.

In addition to arabidopsis, growth defects in *dcl4* mutants have been described in *Solanum lycopersicum* (tomato) (*44*), a dicot separated by ~100 My of evolution from arabidopsis (*45*), and in the moss *Physcomitrium patens (46*), separated by ~400 My of evolution from arabidopsis (*47*). In tomato, *dcl4* mutants are unique in causing leaf necrosis among mutants with defects in biogenesis of endogenous siRNAs (*44*), and in *Physcomitrium patens*, the developmental defects observed in *dcl4* mutants are suppressed by simultaneous inactivation of the *DCL3* gene (*46*). Moss *DCL3* may encode a Dicer-like ribonuclease with functions comprising those of both spermatophyte DCL2 and DCL3, as the DCL2-clade is present only in higher plants (Fig. S2). Strikingly, gene expression profiling showed that the overlap in enriched GO terms among differentially expressed genes in arabidopsis *dcl4-15*, tomato *dcl4 (w3*, Q1277STOP) and *Physcomitrium dcl4-20* compared to wild type centered on defense-related processes (Fig. 1C, Fig. S3A,B). Moreover, differential gene expression between *dcl4* and wild type was substantially, though not fully, restored in *dcl4/dcl3* mutants in *Physcomitrium* (Fig. 1D). We did not test the DCL2-dependence of autoimmune responses in tomato, because its four DCL2 paralogs complicate genetic analyses. Nonetheless, mRNA-seq of an *rdr6* mutant (*wiry*, (*44*)) showed that the transcriptomic changes detected in *dcl4* were not simply a consequence of loss of DCL4-dependent regulatory siRNAs whose biogenesis also requires RDR6 (Fig. S3C). Importantly, the promoters of genes differentially expressed between arabidopsis *dcl4-15* and wild type or *dcl4-15/amiR-DCL2*, and tomato and *Physcomitrium dcl4* and wild type all exhibited significant enrichment of the W-box ((T)TGACY, Y=T/C), a binding site of the defense-related WRKY transcription factors (*48*) (Fig. 1E, see Methods). We conclude that loss of DCL4 in plants leads to autoimmunity characterized by a well-defined WRKY-mediated gene expression program (*49, 50*). Importantly, such autoimmunity depends on a Dicer-like ribonuclease, the spermatophyte-specific DCL2 in arabidopsis, or an ancestral DCL3 in *Physcomitrium*.

## Autoimmune responses require cytoplasmic DCL2

Contrary to other arabidopsis DCLs, no DCL2 isoform contains clear nuclear localization signals (Fig. S4). To clarify whether cytoplasmic or nuclear DCL2 induces autoimmunity, we assessed the penetrance of the autoimmune phenotype in three groups of stable transgenic lines: *dcl4-2t/dcl2-1* expressing FLAG-DCL2^WT^, FLAG-DCL2^NLS^ (with a strong nuclear localization signal) or FLAG-DCL2^NES^ (with a nuclear export signal), all under the control of the endogenous *DCL2* promoter. At similar expression levels, FLAG-DCL2^NLS^ caused significantly lower penetrance of growth phenotypes than FLAG-DCL2^NES^ (Fig. 2A). Because of the low expression levels of DCL2, we used the patterns of endogenous 22-nt siRNAs produced to infer the subcellular localization of the encoded proteins. These analyses verified that DCL2^WT^ produced both endogenous hairpin-derived siRNAs (*IR71*, presumed to be nuclear) and mRNA-derived siRNAs (*SMXL5*, presumed to be cytoplasmic), while DCL2^NLS^ produced mostly hairpin-derived siRNAs, and DCL2^NES^ produced mostly mRNA-derived siRNAs (Fig. 2B). These observations indicate that activation of autoimmunity via DCL2 is particularly dependent on cytoplasmic DCL2.

**Fig. 2.**
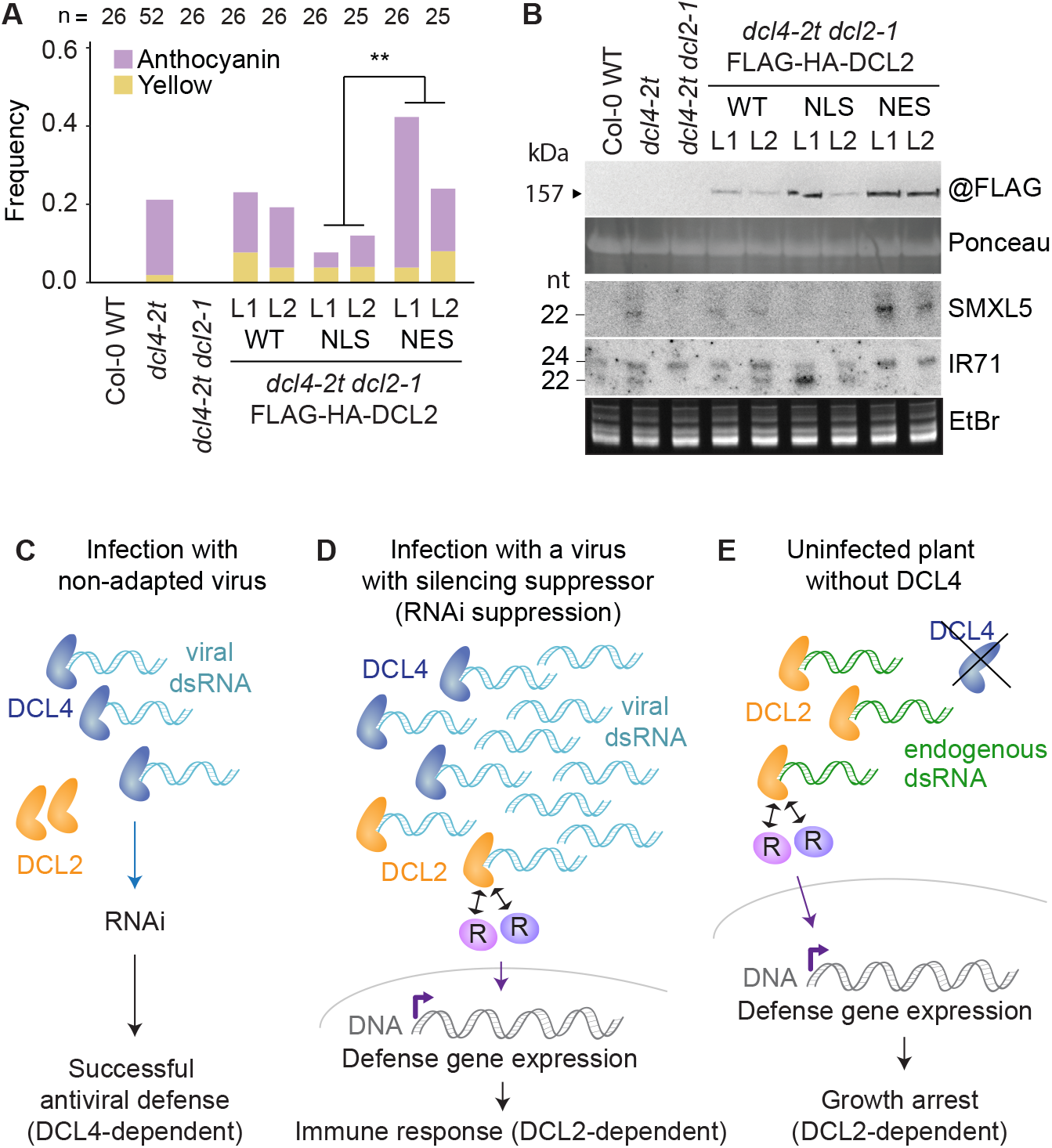
Relevance of cytoplasmic DCL2 for immune activation and a model for a two-layered DCL-mediated innate antiviral immune system. (**A**) Quantification of rosette phenotypes of 28-day old plants of the indicated genotypes, grown under long-day conditions. Two independent transgenic lines (L1, L2) of *dcl4-2t/dcl2-1* expressing FLAG-HA-DCL2 wild type or fused to a C-terminal nuclear localization signal (NLS) or nuclear export signal (NES) were used. Yellow, proportion of plants with visible leaf yellowing; anthocyanin, proportion of plants with visible purple leaf coloring on abaxial sides of leaves. **, significant difference (P < 0.01 by logistic regression, see Methods). (**B**) Molecular analyses of 14-day old sterile-grown seedlings with no apparent phenotypes of the lines used in A. Top, protein blot of total lysates developed using anti-FLAG antibodies. Ponceau staining of the membrane is shown as loading control. Bottom, small RNA blot hybridized to radiolabeled probes complementary to the SMXL5 mRNA and to the endogenous inverted repeat IR71. Hybridizations were carried out consecutively to the same membrane after probe stripping. Bottom panel, EtBr staining of the upper part of the gel as a loading control. (**C-E**) A model for basal antiviral immunity with distinct functions of DCL4 and DCL2. (**C**) A non-adapted virus is eliminated by DCL4-mediated RNAi, without the need for growth arrest and defense gene expression. (**D**) A partially adapted virus capable of RNAi suppression accumulates to levels so high that cytoplasmic DCL2 has access to dsRNA. Cytoplasmic dicing by DCL2 is sensed by R proteins whose activation induces genetic reprogramming leading to growth restriction, sometimes accompanied by programmed cell death, and defense gene expression. (**E**) In the absence of DCL4, DCL2 gains access to endogenous dsRNA, leading to autoimmunity via the same mechanism as in D.

## A new model for DCL-mediated basal antiviral immunity

The results presented so far show that plants share a deeply conserved molecular mechanism to activate innate immune responses in the absence of DCL4. Because the stunted growth phenotype resulting from autoimmunity in arabidopsis *dcl4* mutants is RDR6-dependent (*38, 39*) and is only observed in the absence DCL4 protein, not in plants encoding a total loss-of-function DCL4 point mutant protein that remains bound to dsRNA substrates (*51*), we infer that activation of innate immune responses occur when DCL2, or the ancestral DCL3, gets access to dsRNA, in particular cytoplasmic dsRNA. To explain these observations, we propose a new model for basal antiviral immunity in plants. Rather than acting redundantly to mediate RNAi, DCL4 and DCL2 perform distinct functions in basal antiviral defense (Fig. 2C-D). DCL4 is the major cytoplasmic Dicer activity (*52*), and is dedicated to antiviral RNAi (Fig. 2C). Cytoplasmic DCL2, on the other hand, does not produce siRNAs unless viral dsRNA concentrations reach high levels as a consequence of RNAi inactivation by viral anti-RNAi effectors that constitute a key basis for adaptation to a specific host. In that case, R proteins sense processing of cytoplasmic dsRNA by DCL2 to switch on innate immune responses (Fig. 2D), and thereby provide a second basal layer of antiviral immunity distinct from RNAi. In this model, the absence of DCL4 leads to DCL2-dependent autoimmunity because endogenous cytoplasmic dsRNAs, diced by DCL4 in wild type plants, become substrates of DCL2 (Fig. 2E). Similarly, extreme abundance of dsRNA in mutants of RNA decay factors may saturate DCL4, leading to DCL2-dependent growth repression (*39, 42*). This alternative model for DCL2 function makes several predictions that distinguish it from DCL2’s role in classical RNAi. Most importantly, *R* gene(s) must exist whose inactivation together with *DCL4* abrogates resistance to non-adapted viruses to a degree similar to that observed in *dcl4/dcl2* mutants. The model also predicts that those *R* genes should be identifiable genetically as suppressors of DCL2-dependent autoimmunity in *dcl4* mutants.

## Specific NLRs are required for DCL2-dependent autoimmunity

We used a library of dominant negative *R* genes (*20*) to screen directly for NLRs required for to DCL2-dependent autoimmunity. We reasoned that at least some *R* genes implicated in DCL2-dependent immune activation may show sequence conservation across ecotypes, because the detected molecular cue, cytoplasmic dicing by DCL2, would be invariant. We therefore prioritized arabidopsis *R* genes for screening according to the degree of conservation across 80 ecotypes (*53*). This approach identified the plasma membrane-localized CNL encoded by the *R* gene
*L5* (AT1G12290) (*54*) as a dominant suppressor of *dcl4* (Fig. S6A). We verified that two independent insertion alleles of *L5* clearly, albeit not fully, suppressed *dcl4* growth phenotypes (Fig. 3A). Because the double mutant *dcl4/sgt1b* defective in an Hsp90 co-chaperone proposed to mediate proteolysis of many R proteins (*55, 56*) exhibits stronger, completely penetrant, yet fully DCL2-dependent growth inhibition (Fig. 4B, Supplemental Text 1, Fig. S5), we used this background to further characterize the requirement of *L5*. Knockout of *L5* suppressed the growth phenotype of *dcl4/sgt1b* (Fig. 3B), and the gene expression profile in *dcl4/sgt1b*, with activation of defense-related genes and repression of cell cycle-related genes, was also strongly dependent on *L5* (Fig. 3C). Because the suppression of DCL2-dependent growth phenotypes was not complete, we rescreened the dominant negative *R* gene library, this time focusing on candidates exhibiting gene expression correlated with *L5*, as judged by the ATTED expression database (*57*). This approach identified the TNL *RPP9/RAC1* (AT1G31540) as an additional suppressor of *dcl4* (Fig. S6A). Some alleles of *RPP9* confer resistance to two distinct oomycete pathogens (*58, 59*), a class of pathogen for which *trans*-kingdom RNAi plays a crucial role in determining the outcome of the host-pathogen interaction (*60*). We confirmed that genetic inactivation of *RPP9* suppressed *dcl4/sgt1b* phenotypes (Fig. 3B and D, Fig. S6B-C), and also noted that a combination of *l5* and *rpp9* did not enhance suppression compared to the single mutants (Fig. 3B, Fig. S6B-C), suggesting serial rather than parallel action of L5 and RPP9. Despite the clear suppression of *dcl4/sgt1b*, we noticed some phenotypic variation, particularly among *dcl4/sgt1b/l5* individuals. This included rare cases resembling autoimmune mutants without extensive cell death (Fig. S6B-D)(*49, 61*), thus suggesting involvement of as yet unidentified R proteins (Supplemental Text 2). Notwithstanding this complication, we conclude that at least two R proteins, the CNL L5 and the TNL RPP9, contribute non-redundantly to DCL2-dependent activation of immune responses.

**Fig. 3.**
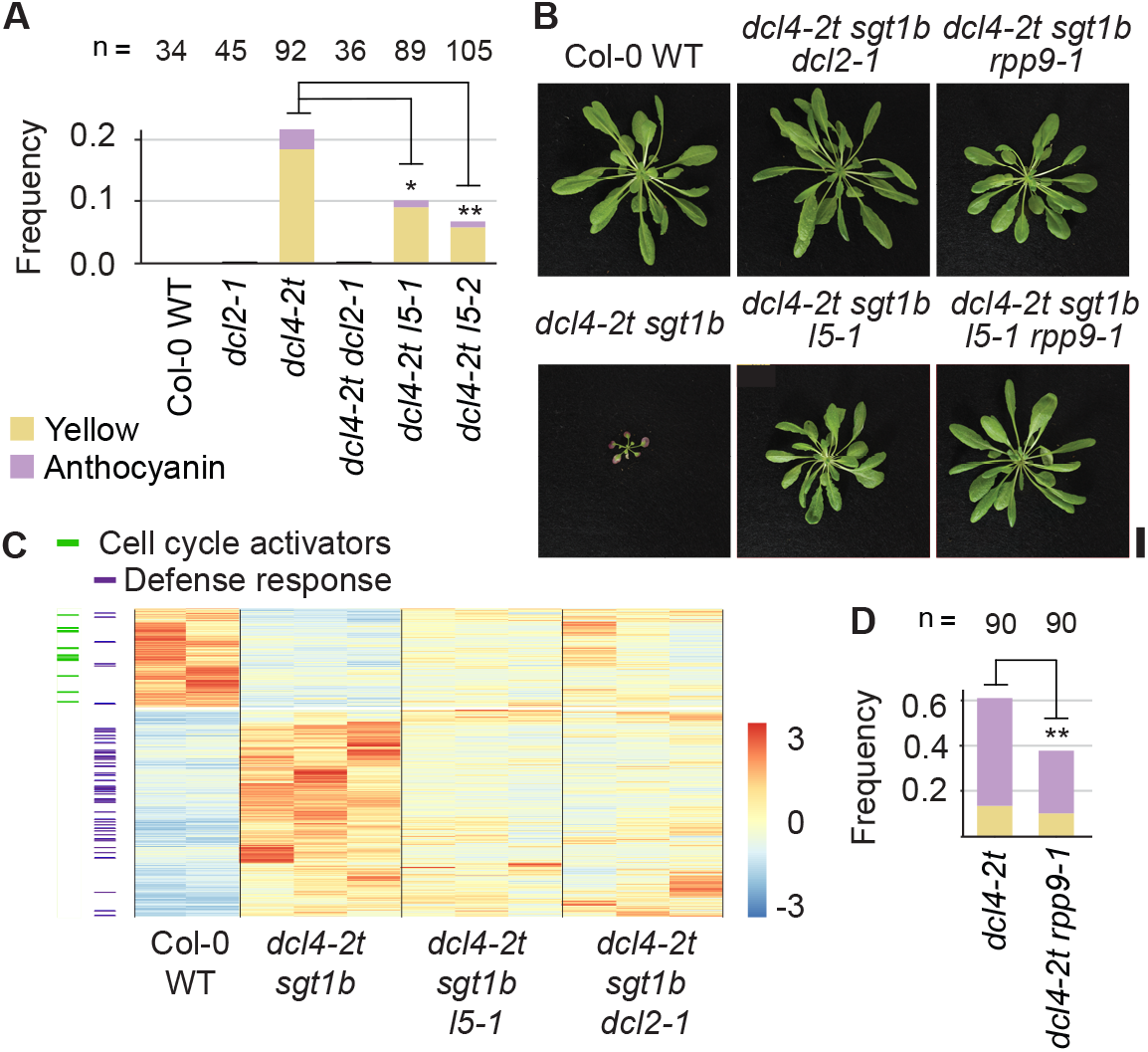
The R proteins L5 and RPP9 are required for DCL2-mediated immune responses. (**A**) Quantification of rosette phenotype occurrence in the indicated genotypes, as in Figure 2A. Asterisks indicate significance of difference compared to *dcl4-2t* (χ^2^ test). ** P < 0.01, * P = 0.05. (**B**) Photographs of representative rosette phenotypes after 25 days of growth under short-day conditions of the indicated genotypes. Scale bar, 2 cm. (**C**) Row-normalized heat map showing genes differentially expressed between the indicated genotypes as determined by mRNA-seq. The column on the left indicates known cell cycle (green) and defense response (purple) genes. (**D**) Quantification of rosette phenotype occurrence in the indicated genotypes, as in Figure 2A. **, P < 0.01 (χ^2^ test).

## Basal antiviral resistance requires R proteins functionally linked to DCL2

We finally tested whether L5 and RPP9 mediate basal antiviral immunity in the absence of RNAi. This should be detectable as loss of resistance to a virus deprived of its anti-RNAi effector only when *l5* and *rpp9* mutations are combined with *dcl4*, but not in *l5* and *rpp9* single mutants. Furthermore, the expected R protein-dependence of the antiviral DCL2 function predicts that inactivation of DCL2 and R proteins should not show additive effects on loss of antiviral immunity. We chose the positive strand RNA virus turnip crinkle virus without its anti-RNAi effector P38 (TCVΔP38) (*62*) to test these predictions. Inactivation of *L5* had no effect on resistance to TCVΔP38, but caused mild loss of antiviral resistance in combination with *dcl4*, comparable to *rdr6* or to *dcl2* in the heterozygous state as shown in the accompanying paper (Fig. 4A-B) (). Accumulation of 22-nt viral siRNAs was not affected by mutation of *L5* (Fig. 4A). Furthermore, inactivation of *RPP9*, or combined inactivation of *RPP9* and *L5*, caused stronger loss of resistance to TCVΔP38 in combination with *dcl4*, and neither *l5* nor *rpp9* mutants enhanced the susceptibility of *dcl4/dcl2* mutants (Fig. 4B). These results demonstrate that NLRs functionally linked to DCL2 constitute a crucial layer of basal antiviral resistance when RNAi has been abrogated by mutation of *DCL4*.

**Fig. 4.**
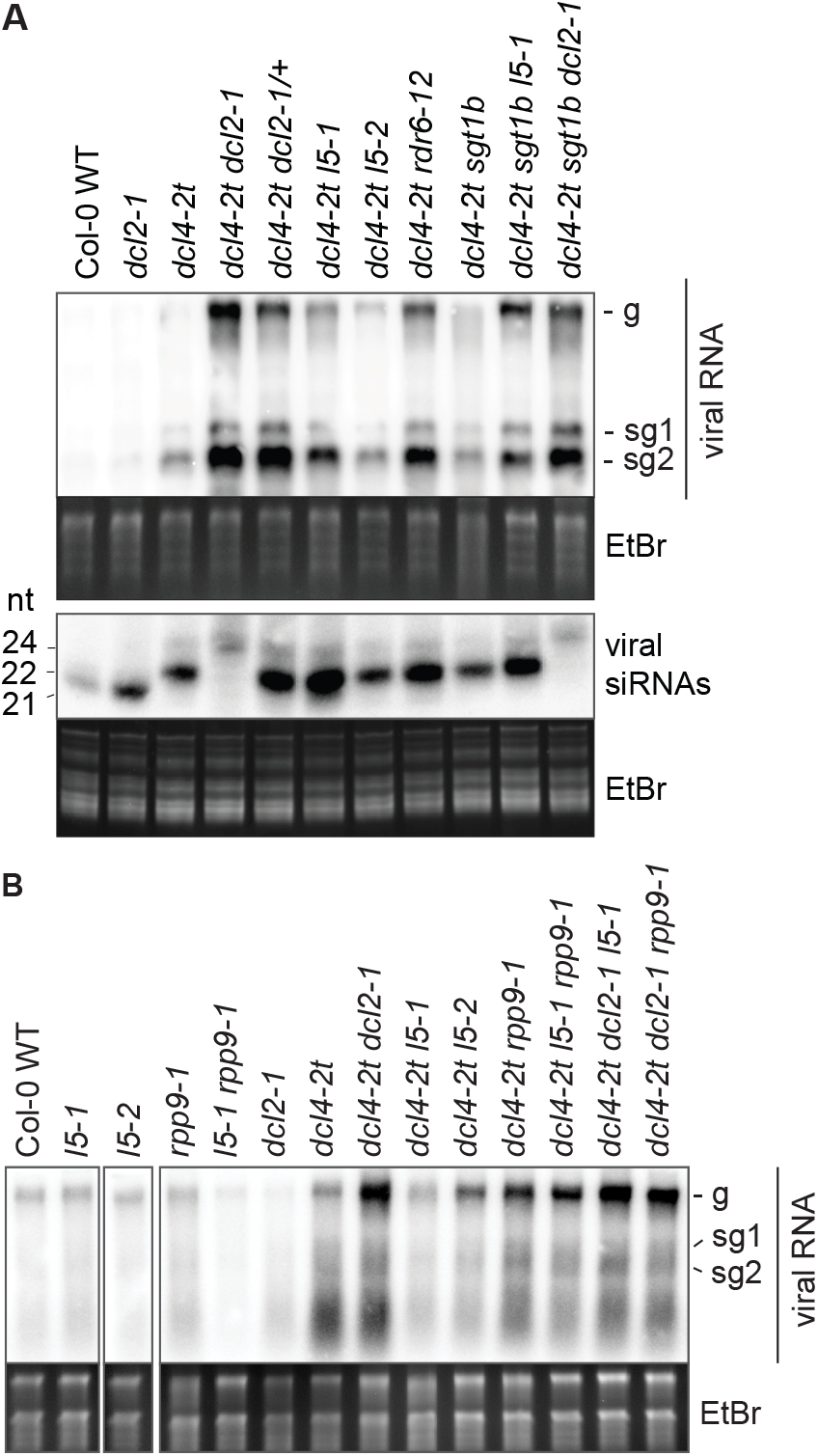
RNAi and DCL2/R proteins constitute distinct layers of immunity against TCV P38. (**A**) Top, accumulation of viral gRNA in TCVΔP38-infected leaves at 5 days post-inoculation, assessed by RNA blot. Bottom, small RNA blot showing TCV-derived siRNAs in the same leaves. (**B**) Same as in A (top) for a different set of genotypes. g, genomic RNA; sg, subgenomic RNAs. EtBr staining is shown as loading control in all cases.

## DISCUSSION

Our study provides strong genetic evidence to support a new conceptual framework for basal antiviral resistance in plants. The two antiviral DICER-LIKE ribonucleases mediate distinct layers of antiviral activity, such that DCL4 initiates the primary defense layer, RNAi, while DCL2 mediates NLR activation when RNAi is defeated and cytoplasmic dsRNA concentrations reach high levels. It is formally possible that ectopic DCL2-dependent siRNAs in *dcl4* mutants coincidentally silence mRNAs encoding host proteins guarded by L5 and/or RPP9, leading to their activation. Such a scenario is strongly disfavored by two observations. First, the conservation of DCL-dependent autoimmunity triggered by inactivation of *DCL4* between moss and arabidopsis makes fortuitous silencing of genes conditioning R protein activation unlikely. Second, the accompanying manuscript shows that plants with reduced DCL2 dosage or with specific helicase mutations in DCL2 did not show substantially lower levels of endogenous 22-nt siRNAs, yet rescued auto-immune phenotypes (). Thus, our study uncovers a fundamental element of basal antiviral immunity, likely to be conserved throughout the plant kingdom. Sensing of cytoplasmic dsRNA is also a crucial element of innate antiviral immunity in several different animal species (*27*). With our discovery of a function of DCL2 and the NLRs L5 and RPP9 in dsRNA sensing, it now appears that dsRNA sensing through Dicers, Dicer-related helicases and NLRs is a recurrent theme in eukaryotic innate immunity. This emerging unifying principle deserves further consideration. For example, mammalian cytoplasmic dsRNA sensors include two RLRs (*27*) with DExD/H-box helicase domains whose closest relatives are those of Dicers. In *C. elegans*, a RIG-I-like protein, Dicer-Related Helicase DRH1, plays roles in antiviral RNAi (*63, 64*), but recent studies show that it also mediates defense-related transcriptional re-programming in response to viral RNA (*65*). A dual role in RNAi and dsRNA sensing may also apply to *Drosophila* Dicer-2 (Dcr-2) that is required for antiviral RNAi via the ARGONAUTE protein Ago-2 (*66–68*). Indeed, while *dcr-2* and *ago-2* mutants share defects in antiviral RNAi (*67*), *dcr-2* mutant flies also have defects in virus-induced defense gene expression (*69*), indicating additional roles of Dcr-2 in immune signaling.

In contrast to mammalian RLR-signaling, dsRNA-dependent activation of plant immune responses via DCL2 involves NLRs, whose activation by oligomerization is driven by ATP binding (*9, 70, 71*). Remarkably, however, examples of dsRNA perception relayed via NLRs have also been found in animals whose R-protein equivalents are Nucleotide Oligomerization Domain (NOD)-like receptors that form inflammasomes upon oligomerization (*72*). A mammalian DExD/H helicase, Dhx9, senses rotaviral RNA and activates innate immune responses via the NOD-like receptor NLRP9b (*73*), and in humans, NLRP1 has a recently evolved function to sense dsRNA (*74*). Finally, some plant TNL-based resistosomes use dsRNA/dsDNA substrates to synthesize 2’,3’-cAMP/cGMP with cell death-promoting signaling activity, independently of their NADase activity (*75*). It appears, therefore, that Dicer-related helicases, sometimes Dicers themselves, and NLRs are at the core of cytoplasmic dsRNA sensing in eukaryotes. In some cases, the helicase-mediated perception of dsRNA is relayed via formation of inflammasome-type oligomers of nucleotide-binding signaling proteins, in other cases signaling proceeds directly from sensing helicases, or may involve inflammasomes/resistosomes directly.

It is unclear whether L5 and RPP9 participate directly in sensing or whether one or both of them transduce the signals from the sensor, as has been established for some so-called helper-NLRs (*76, 77*). Finally, we stress that the cytoplasmic DCL-dependent dsRNA sensing system we have discovered is unlikely to be the only mode of dsRNA sensing in plants. First, a DCL-independent function of exogenously applied dsRNA as a PAMP in arabidopsis has been described (*78*), implying the existence of cell-surface or, as in mammals, endocytic dsRNA sensors (*26*). Second, our transcriptome analyses of *dcl4/dcl2* in arabidopsis and of *dcl4/dcl3* in *Physcomitrium*, did not show complete abrogation of the immunity gene expression signature of *dcl4* mutants. Thus, similar to metazoans (*26, 79, 80*), additional cytoplasmic dsRNA sensors may exist, a possibility supported by the recent demonstration of broad antiviral activity of the cytoplasmic dsRNA-binding protein DRB2 in arabidopsis (*81*).

## ACKNOWLEDGMENTS

Detlef Weigel is thanked for support and allocation of laboratory resources to the study of R protein conservation across Arabidopsis accessions. Theo Bölsterli, Rene Hvidberg and their teams are thanked for plant care. Simon Bressendorff and Jacob Kanne are thanked for assistance with growth of *Physcomitrium patens*. The Nottingham Arabidopsis Stock Centre (NASC) is thanked for providing Arabidopsis T-DNA insertion lines, and the International Moss Stock Center (IMSC), Freiburg, Germany, is thanked for providing *Physcomitrium patens dcl4* and *dcl3* mutants. Yuval Eshed is thanked for providing seeds of tomato wild type, *dcl4* and *rdr6*. Mahmot Tör (*edm1*) and Jane Parker (*sgt1b-1*) are thanked for providing seeds of Arabidopsis *sgt1b* mutants. Detlef Weigel, Christophe Ritzenthaler, Simon Bressendorff, Diego López-Márquez, Emilie Oksbjerg and Dexter Adams are thanked for critical reading of the manuscript. This work was supported by grants to P.B. from the Independent Research Fund Denmark (Sapere Aude Grant 12-133793), Villum Fonden (Project 13397), and the European Research Council (PATHORISC, ERC-2016-CoG 726417).

## AUTHOR CONTRIBUTIONS

CPSN acquired and analyzed all high-throughput sequencing data, identified *L5* by reverse genetic screening and constructed and analyzed genetic backgrounds containing *l5* mutant alleles, characterized *dcl4/sgt1b* double mutants, LH constructed and characterized *dcl4-15/*amiR-*DCL2* lines, isolated *RPP9* by reverse genetic screening and constructed and analyzed all genetic backgrounds containing *rpp9* mutant alleles, LA-H acquired the first evidence for defense activation in *dcl4* mutants, constructed double and triple mutants with *dcl4-2t* and *eds1, sid2, nahG, sgt1b*, initiated screening of dominant negative *R* genes, and participated in discussions with PB leading to formulation of the model for DCL2 function proposed here, DK analyzed R protein conservation across Arabidopsis accessions and related crucifer species, MP provided the library of dominant negative *R* genes, PB conceived the project, designed experiments, supervised work and wrote the manuscript with input from all authors.

## METHODS

### Plant materials and growth conditions

Unless otherwise stated, plants were grown in soil (Plugg/Såjord (seed compost), SW Horto) under a long day cycle (day: 16 hours light, 130 μmol m^-2^ s^-1^, 21°C; night: 8 hours darkness, 18°C), in Percival growth chambers equipped with Philips Master TL-D 36W/840 and 18W/840 bulbs. *A. thaliana* seeds were surface sterilized before stratification and germination. Surface sterilization was performed by treatment with 70% ethanol for 2 minutes, then in 1.5 % NaOCl, 0.05 % Tween-20 for 10 minutes, followed by two washes in deionized water. After surface sterilization seeds were stratified by incubation in the darkness at 4°C for 3 days. Seeds were then germinated on either soil (Plugg/Såjord (seed compost), SW Horto) or on Murashige-Skoog (MS) agar plates (4.4 g/L MS salt mixture, 10 g/L sucrose, 8 g/L agar, pH 5.7).

*P. patens* lines were grown on BCDAT media (*82*) solidified with 8 g/L agar and overlaid with cellophane discs (AA packaging), for 3 weeks under constant temperature (21 °C) and a light intensity of 70 μmol m^-2^ s^-1^. Tissue was harvested in liquid nitrogen.

*S. lycopersicum* lines were germinated in soil (Plugg/Såjord (seed compost), SW Horto) and grown under long day conditions in the greenhouse. Leaf tissue was harvested in liquid nitrogen at 48 days post germination. At this point in time, anthocyanin accumulation was visible on the abaxial sides of petioles in *dcl4*, but not in *rdr6* or wild type. Apart from the signs of anthocyanin accumulation, the leaves of *dcl4* did not look different from those of *rdr6*.

### Plant genotyping

All T-DNA lines were genotyped using PCR with 2 different primer sets. One primer set to detect the wild type allele with forward and reverse primers on opposite sides of the T-DNA. The other primer set detects the T-DNA insertion allele, and uses either the forward or reverse primer from the first set, together with a primer inside the left border of the T-DNA. For point mutants, derived Cleaved Amplified Polymorphic Sequence (dCAPS) markers (*83*) were designed to allow genotyping by restriction digest following PCR. The Col-0 *sgt1b* (*edm1*) mutant is a large 35 kb deletion (*84*). This mutant was genotyped using primers inside the *SGT1b* gene (LA288/LA290), so that only the presence of the wild type allele could be demonstrated directly. Repeated absence of reactivity was taken as proof of homozygosity of the *sgt1b* (*edm1*) mutant allele. All primer sets and, if appropriate, cognate restriction enzymes are listed in Table S3.

### Quantification of incompletely penetrant phenotypes

Seeds were surface sterilized, plated on MS medium and stratified at 4°C for 3 days. The plates were then incubated at constant temperature (22°C) with a 16-hour light/8-hour darkness cycle. After 2 weeks, seedlings were transferred to soil, and after another 2 weeks the phenotype was scored. For the statistical analysis, we categorized plants showing yellowing and anthocyanin accumulation as one category (symptomatic). To compare the fraction of healthy plants and plants showing autoimmune symptoms between individual lines, we used a standard chi-squared test in R. To compare the fraction of healthy and symptomatic plants between groups of lines, as for example in the comparison of the two DCL2-NLS and DCL2-NES lines, we used logistic regression in R using the gmodels package. To assess the phenotype of plants with a *dcl4-2t/sgt1b* double mutant background, the seeds were germinated on soil and grown under short day conditions (8h light/16h darkness). Phenotypes were assessed at a point in time at which all *dcl4-2t/sgt1b* individuals showed a clear autoimmune phenotype, typically 4-6 weeks after germination and before bolting.

### RNA extraction

RNA was extracted by mixing 150-200 mg of plant tissue ground under liquid nitrogen with 1 ml of TRI reagent (Sigma). 200 μl of chloroform was added and after vortexing, samples were centrifuged at minimum 16000x *g* for 15 minutes at 4°C. The aqueous phase was transferred to a new tube and mixed with isopropanol in a 1:1 ratio, after which the samples were incubated at −20°C for 1 hour or over night. Precipitated RNA was pelleted by centrifugation at 16000x *g* for 15 minutes. Pellets were washed twice with ice-cold 70 % ethanol and were then dissolved in 300 μl of milliQ water. To precipitate polysaccharides, we then added 10 μl of 3M NaOAc, pH 5.2, 30 μl ice cold 96% ethanol and incubated samples on ice for 30 minutes. Polysaccharides were pelleted by centrifugation and supernatants were transferred to new tubes. 30 μl of 3M NaOAc pH 5.2, and 750 μl of 96 % ethanol were then added, and samples were incubated at −20°C for 1 hour or overnight. Precipitated RNA was pelleted by centrifugation and washed twice with 70 % ethanol before it was dissolved in Milli-Q water for sequencing or quantitative RT-PCR analysis, or in 50 % formamide for northern blot analysis.

### Quantitative RT-PCR

#### cDNA synthesis

To produce cDNA, 3 μg of total RNA were mixed with 1 U DNAse I, 0.2 U Ribolock and 1x DNAseI buffer (Thermo Scientific), and incubated at 37°C for 30 minutes. DNaseI was inactivated by addition of 1 μl of 50 mM EDTA and incubation at 65°C for 10 minutes. 4 μl of the DNAseI reaction was mixed with 1 μl of oligo(dT)_18_ 0.5 μg/μl (Thermo Scientific), and incubated at 65°C for 5 minutes. The 5 μl were then added to 16 μl of an RT mastermix (1U RTase, 0.5 U Ribolock, 4 mM dNTP mix, 1x RT buffer (Thermo Scientific)), and incubated at 42°C for 1 hour, followed by a 10minute incubation at 70°C to inactivate the RTase.

#### Quantitative PCR

0.5 μl of cDNA was mixed with appropriate qPCR primers (see Table S3) and Maxima SYBR Green qPCR Master Mix (Thermo Scientific). The qPCR reactions were run in technical triplicates on a Bio-Rad CFX ConnectTM thermal cycler, and the obtained Ct values were used to calculate relative expression using the ΔΔCt method (*85*).

### mRNA-seq, differential gene expression analysis and GO term enrichment

As a general rule, sequencing experiments used biological triplicates in which independent pools of biological material (three for each mutant background) were grown and harvested separately, but in parallel, under the same conditions. The only exceptions to this were the tomato mRNA-seq analysis that used duplicates, and the Col-0 background in the mRNA-seq experiment in which Col-0, *dcl4/sgt1b, dcl4/sgt1b/dcl2-1* and *dcl4/sgt1b/l5* backgrounds were compared. In this case, one of the Col-0 samples failed, and only two replicates were used in the analysis.

mRNA was isolated from total RNA using the NEBNext PolyA mRNA Magnetic Isolation Module. The libraries were prepared with the NEBNext Ultra RNA Library Prep Kit for Illumina (#E7530S) using Agencourt AMPure Beads for the cleaning steps. Quality of libraries and input RNA was assessed using an Agilent 2100 Bioanalyzer. Libraries were sequenced by Novogene Bioinformatics Technology Co., Ltd on an Illumina HiSeq 2500 platform. For the tomato and moss samples the library preparations were also done by Novogene Bioinformatics Technology Co., Ltd using 2 μg of total RNA as input. FASTQ files were trimmed using Cutadapt (*86*) and mapped using STAR (*87*). Genes were counted using featureCounts 1.6.3 (*88*), and imported into R where data analysis and identification of differentially expressed genes was done using DESeq2 (*89*). Gene Ontology Analysis was done with g:Profiler (*90*), using genes with more than 10 counts in the whole dataset as background.

### Motif enrichment analysis

To analyze the enrichment of the W-box in the promoters of upregulated genes in *P. patens*, tomato and *A. thaliana dcl4* mutants, we first performed *de novo* motif discovery using HOMER2 (*91*). We noticed that in several of the sets, HOMER2 identified different variants of the W-box as being significantly enriched. We chose the W-box-like motif identified by HOMER2 in the comparison of moss *dcl4-20* vs WT for in-depth statistical analysis of enrichment, because of the similarity of this motif (TTGACY(D), Y=C/T, D=T/A/G) to the canonical W-box ((T)TGACY). We then used FIMO (*92*) to assess the significance of the enrichment of the W-box found in HOMER2 in the different sets of differentially expressed genes.

### Cloning

The DCL2 WT/NES/NLS constructs were created by USER cloning. We used the SV40 T-antigen NLS (PKKKRKVEDP) and the HIV rev NES (LQLPPLERLTLD). These are both from mammalian viruses, but retain NLS and NES activities, respectively, in protoplasts (*93*). USER primers with appropriate overhangs were designed so that DCL2 could be amplified in 2-3 fragments. The first fragment was amplified using CSN90/CSN79, and the forward primer CSN90 is complementary to the promoter region of DCL2 while the reverse primer CSN79 is complementary to the CDS of DCL2 creating a fragment of 4739 bp. This fragment was the same for all three constructs. The plasmid containing Pro(DCL2):2xFLAG-2xHA-DCL2^WT^:ter(35S) was used as template for this reaction so that we included the N-terminal 2xFLAG-2xHA tag.

For the wildtype construct we only used 2 fragments, and the second fragment was amplified using CSN78/CSN91 and genomic DNA as template in order to retain the native DCL2 terminator. CSN78 has a USER overhang complementary to CSN79 so that it can be ligated to fragment 1, while CSN91 anneals to the DCL2 terminator (355 bp downstream of the stop codon).

The DCL2^NLS^- and DCL2^NES^-encoding DNA sequences were cloned in 3 fragments. The second fragment used a forward primer in the CDS of DCL2 matching fragment 1 and a reverse primer at the STOP codon, which contains either an NES or an NES in its USER overhang. CSN78/CSN66 was used for the NLS construct and CSN78/CSN69 was used for the NES construct. For the third fragment we used CSN67/CSN91 for the NLS construct and CSN70/CSN91 for the NES construct, and genomic DNA as template. This produces overhangs that match the ones in fragment 2 and the USER cassette. In total the ligation of these fragments results in constructs containing the DCL2 promoter, the DCL2 gene with an N-terminal 2xFLAG-2xHA tag, and the DCL2 terminator. The only difference between the three constructs is the presence or absence of an NLS/NES encoded at the C-terminus. The sequences of all primers used are listed in Table S3.

### Plant transformation and selection of transgenic plants

Plants were transformed with the appropriate vectors using floral dip transformation (*94*). *Agrobacterium tumefaciens* strain GV3101 was transformed with the relevant binary vector, selected on appropriate antibiotics and grown in liquid LB to an OD_600_ of 0.8-1.2. The culture was then centrifuged and the bacterial pellet was resuspended in infiltration medium (5 % sucrose, 0.05 Silwet L-77 (Lehle seeds)), to an OD_600_ of 0.8. Flowering plants were then submerged in the bacterial broth for about 1 minute, and left to set seeds.

T1 seeds transformed with constructs in pB7m34GW and in pLIFE41 were sown on soil and sprayed with glufosinate ammonium at 14,16 and 18 days after germination. Seeds of a total of 30 resistant T1 plants were harvested and kept for further selection.

T2 seeds were then grown on MS agar plates containing 7.5 mg/L glufosinate ammonium. The number of sensitive and resistant seedlings were counted to identify lines having a ratio of sensitive plants of 25 %, indicating that these contained one copy of the transgene. 12 plants from each of 3-4 lines were transferred to soil and harvested after ripening. T3 seeds were again sown on MS agar containing appropriate antibiotic to identify seeds batches with 100 % resistant plants indicating that these were homozygous for the transgene. Homozygous T3 seed batches from 3-4 lines containing a single transgene were then used for further experiments.

### Protein blotting

The aerial parts of seedlings were fast frozen in liquid nitrogen and ground to a fine powder. The powder was mixed with IP buffer (50 mM Tris-HCl pH 7.5, 150 mM NaCl, 10 % glycerol, 5 mM MgCl2, 0.1 % Nonidet NP-40) in the ratio of three ml IP buffer per g of tissue. Samples were then centrifuged at 16000 x *g* at 4°C to pellet cell debris. Supernatant was mixed with LDS loading buffer (70 mM Tris-HCl, pH 6.8, 10 % glycerol, 1 % lithium dodecyl sulfate (LDS), 0.01 % bromophenol blue), boiled at 85 °C for 5 minutes. Samples were loaded on a precast 4-20 % Criterion gradient gels (Biorad), which was subjected to SDS PAGE and blotted on a nitrocellulose membrane (Amersham Protran Premium 0.45 μm, GE Healthcare). The blotted membrane was blocked in 5% skimmed milk for 1 hour. After blocking the membrane was incubated with Monoclonal anti-FLAG HRP antibody (Sigma, F3165) overnight at 4°C. The following day the membrane was washed 3 times 5 minutes in PBS-T, after which the HRP-conjugated anti-FLAG antibody signal was detected using enhanced chemiluminescence.

### Small RNA blotting

RNA was mixed with small RNA loading dye (20 mM HEPES pH 7.8, 1 mM EDTA, 50 % formamide, 3 % glycerol and 0.01 % bromophenol blue), denatured at 65 °C for 10 minutes, loaded on a 18% acrylamide:bis 19:1, 7 % urea, 0.5X TBE gel, and run in 0.5X TBE buffer until the bromophenol blue had reached the bottom of the gel. After separation the gel was blotted onto an Amersham Hybond-NX nylon membrane (GE Healthcare Life Sciences), after which RNA was crosslinked using carbodiimide-mediated crosslinking (*95*). Membranes were then rinsed in deionized water and prehybridized with PerfectHybTM Plus Hybridization Buffer (Sigma) at 42°C for 20-60 minutes. The appropriate probe was added and left to hybridize overnight at 42°C. The following day the membrane was washed 3 times 20 minutes in 2xSSC (0.3 M NaCl, 30 mM sodium citrate) 2 % SDS at 42-50 °C and exposed on a phosphor screen. After satisfactory signal was obtained membranes were stripped by washing 3 times with boiling 0.1 % SDS, in order to enable rehybridization with other probes. For IR71, SMXL5 and viral siRNA probes, we PCR amplified the appropriate fragments from either plant genomic DNA or the TC VΔP38-containing vector T1D1. The probes were then produced by using the Prime-A-Gene labeling kit (Promega) according to the manufacturer’s instructions.

### RNA gel blotting (viral genomic RNA)

RNA was mixed with RNA loading dye (20 mM KOH buffered HEPES pH 7.8, 60 % formamide, 20 % formaldehyde, 0.1X 6x DNA loading dye (Thermo Scientific) (1 vol sample to 3 vol loading dye), 5 % ethidium bromide), denatured for 10 minutes at 65°C before it was separated on a denaturing 1 % agarose formaldehyde gel (1% agarose, 20 mM HEPES-KOH pH 7.8, 6% formaldehyde, 1 mM EDTA). The separated RNA was blotted onto a Amersham Hybond-NX nylon membrane (GE Healthcare Life Sciences) membrane by capillary transfer, and crosslinked to the membrane either by chemical crosslinking (see sRNA northern) or by UV crosslinking (254 nm) using a Stratalinker (Stratagene).

### Infections with TCVΔP38

Viral RNA was prepared by *in vitro* transcription of a pT1D1 plasmid template linearized by XbaI digestion. *In vitro* transcription was performed with the RibomaxTM Large Scale RNA production kit (Promega) in the presence of Ribo m^7^G Cap analog (Jena Bioscience), using the manufacturer’s guidelines. *In vitro* transcribed RNA was purified by phenol:chloroform extraction and dissolved in MQ water.

Surface sterilized seeds were stratified for 3 days after which they were germinated on soil. Three weeks after germination, leaves were inoculated with *in vitro* transcribed TCVΔP38 RNA according to the protocol used in (*62*). Briefly, leaves were gently rubbed with silicon carbide to create lesions for viral RNA entry and 20 μl of 10 ng/μl TCVΔP38 RNA was rubbed on the leaf. We infected three leaves per plant and infected 6 plants per genotype. Five days after inoculation, infected leaves were harvested in liquid nitrogen.

### Ranking of Arabidopsis *R* gene conservation across accessions

To estimate sequence conservation, 163 *NLR* genes were ranked based on their conservation values, defined as the fraction of coding sequence length that had non-zero read coverage. Very short reads from 80 A. thaliana accessions (*96*) were trimmed to equal lengths (36bp) and mapped to the reference Col-0 accession (TAIR10 assembly), allowing one mismatch and zero gaps. For each accession, coding sequences of the reference gene were scored for the presence of reads, and fractions of these sequences where at least one read had been mapped were calculated. Conservation value for each gene was calculated as the average of these fractions across the 80 accessions. Details of the mapping and the general approach have previously been published (*53, 97*). The full list of Arabidopsis *R* genes ranked according to conservation across the 80 Arabidopsis accessions is given in Table S1.

### Reverse genetic screening for dominant negative *R* gene suppressors of *dcl4-2t*

The reverse genetic screening was done by direct transformation of dominant negative *R* gene versions into *dcl4-2t* using the published library of dominant negative *R* genes (*20*). Transgenic lines were selected for phosphinothricin resistance on agar plates, transferred to soil at 12-14 days post germination. Phenotypes in the first transgenic generation were inspected relative to a control transformation of *dcl4-2t* with the empty pLIFE41 vector conferring phosphinothricin resistance. The screen was carried out in two stages. In the first, focused on CNLs, dominant negative versions of following R genes were transformed into *dcl4-2t:* AT1G12290, AT5G04720, AT1G12280, AT1G52660, AT3G14460, AT1G50180, AT3G50950, AT4G33300, AT3G14470, AT1G10920, AT5G66910, AT1G53350, AT1G59620. Among those genes, only the dominant negative AT1G12290 (*L5*) version gave rise to appreciably lower frequencies of growth inhibition phenotypes of *dcl4-2t* in the first transgenic generation. In the second, and more broadly focused screen that included both TNLs and CNLs, we screened the following dominant negative R genes: AT1G17600, AT5G22690, AT1G17610, AT3G04220, AT5G18370, AT5G40090, AT5G66900, AT5G40100, AT1G63740, AT5G46450, AT1G72950, AT5G17680, AT5G46470, AT5G18360, AT1G27170, AT2G17050, AT5G38850, AT1G63750, AT1G33560, AT5G11250, AT4G12010, AT1G31540, AT1G72910. Among those genes, the dominant negative version of AT1G31540 (*RPP9*) gave the clearest phenotypic rescue in the first transgenic generation, and was selected for further characterization. Given the noisy nature of the phenotype used for screening, we cannot exclude weak effects of some of the other dominant negative R genes screened on the growth phenotype of *dcl4-2t*.

### Phylogenetic analysis of DCL protein sequences

To construct a phylogenetic tree of land plant DCL proteins, a multiple sequence alignment using Clustal-Ω (*98*) was conducted based on protein sequences obtained from TAIR (https://www.arabidopsis.org/), phytozome (https://phytozome-next.jgi.doe.gov/), and Ginkgo DB (https://ginkgo.zju.edu.cn/genome/). Evolutionary analyses were conducted in MEGA7 (*99*). The phylogenetic tree was generated from the alignment using the neighbor-joining method (*100*). Values of the bootstrap test (*101*) were inferred from 1000 replicates. Branches corresponding to partitions reproduced in <50% of the bootstrap replicates were collapsed. The length of the branches to represent the evolutionary distance in number of amino acid substitutions per site were computed using the Poisson correction method (*102*).

### Data and code availability

This study did not generate new software. mRNA-and sRNA sequencing data sets generated in this study were submitted to the European Nucleotide Archive under accession number PRJEB52819.

## SUPPLEMENTAL INFORMATION TITLES AND LEGENDS

**Fig. S1. The *dcl4-15* knockout mutant constitutively expresses the defense marker genes *PR1* and *SAG13***

Quantitative RT-PCR analysis of relative accumulation of *PR1* and *SAG13* mRNA in 16-day-old seedlings of L*er* WT and *dcl4-15*. cDNA inputs were normalized to *ACTIN2* expression. y-axis shows fold change of expression relative to wild type.

**Fig. S2. Phylogeny of plant DCL proteins**

(**A**) Phylogenetic relations of land plants highlighting the individuals from the major phylogenetic groups from which DCL protein sequences were retrieved. (**B**) Phylogenetic tree of land plant DCL proteins based on multiple sequence alignment produced by Clustal-Ω (see Methods). The length of the branches represents the evolutionary distance in numbers of amino acid substitutions per site. While DCL1, DCL3 and DCL4 are found in all land plants, DCL2 is only found in spermatophytes.

**Fig. S3. Enrichment of functionally related genes in sets of genes upregulated in *Solanum lycopersicum* and *Physcomitrium patens dcl4* mutants**

(**A**) Top 20 gene ontology (GO) terms enriched in genes significantly upregulated in *S. lycopersicum dcl4* compared to wild type, as determined by mRNA-seq. (**B**) Top 20 gene ontology (GO) terms enriched in genes significantly upregulated in *P. patens dcl4-20* compared to wild type, as determined by mRNA-seq. (**C**) Row-normalized heat map showing expression in *S. lycopersicum* wild type, *dcl4* and *rdr6*, of genes differentially expressed between *dcl4* and wild ^t^yp^e^.

**Fig. S4. Presence of predicted nuclear localization signals in Arabidopsis DCL proteins**

Schematic representation of Arabidopsis DCL1-4 proteins. The positions of predicted nuclear localization signals (NLSs), as determined with the NLS prediction software NLS mapper, are highlighted in red. Please notice that *DCL4* is transcribed from two alternative transcription start sites of which only the minor, long form contains an NLS. The NLSs in DCL1, DCL3 and the long form of DCL4 have been described previously (*33, 52, 103*).

**Fig. S5. Visible effects of immune signaling and *sgt1b* mutants on leaf phenotypes of *dcl4* mutants**

(**A**) Frequency of plants showing either leaf yellowing (yellow), visible anthocyanin accumulation (anthocyanin), or both (yellow + anthocyanin) in the indicated genotypes (all in the Col-0 ecotype). 28-day old plants grown under long-day conditions, transferred from MS-agar to soil at day 11 after germination. (**B**) Rosette phenotypes of *dcl4-2t/sgt1b* and *dcl4-2t/sgt1b/dcl2-1*. 49 days of growth, short day, germinated on soil. Size bar, 2 cm. (**C**) Representative photographs of 3-week old rosettes of the indicated genotypes in the L*er* ecotype. As in accession Col-0, the penetrance of the rosette phenotype of *dcl4-15/sgt1b-1* was 100%. Size bar, 1 cm.

**Fig. S6. Genetic interactions between *DCL4, L5* and *RPP9***

(**A**) Quantification of rosette phenotypes indicating the fraction of plants showing visible leaf yellowing (yellow) or anthocyanin accumulation (anthocyanin). DN, dominant negative. The experiments shown use one line (out of many generated) of *dcl4-2t/L5-DN* (left) and of *dcl4-2t/RPP9-DN* (right) propagated to homozygosity for the transgene encoding the dominant negative R protein. Asterisks indicate significance of difference to *dcl4-2t* (χ^2^ test), * P < 0.05, *** P < 0.001. (**B**) Representative individuals exemplifying the degree and variability of suppression of the *dcl4* growth phenotype by additional mutation in the *R* genes *L5* and *RPP9* and their combination, compared to that of *DCL2* mutation. Numbers on the lower-right corner of each photograph indicate the category assigned to each plant as described in C. The two photographs marked with asterisks on the right-upper corner are magnified in D. Scale bar, 2 cm. (**C**) Quantification of rosette phenotypes of the plants photographed in B and additional individuals of each genotype (see n numbers), according to categories 1-4 as described. (**D**) Magnification of the photographs marked with asterisks in panel B, to highlight the autoimmune-like phenotype with severely stunted growth but no visible leaf chlorosis exhibited by a small subset (< 5%) of individuals in *dcl4-2t/sgt1b/l5* populations (category 4 in B and C). Such phenotypes were not observed in *dcl4/sgt1b* populations in either accession Col-0 or L*er*. Size bars, 1 cm.

**Table S1. List of Arabidopsis thaliana *R* genes ranked according to conservation across 80 accessions**

**Table S2. Primers and other oligonucleotides used in the study**

## SUPPLEMENTAL TEXT 1

**Mutation of the Hsp90 co-chaperone SGT1b enhances DCL2-dependent autoimmunity**

We introduced a series of genetic backgrounds with defective *R* gene-dependent immune activation, and/or accumulation of the immune signaling molecule salicylic acid ((*eds1* (*104*), *sid2, sid2/eds1*, overexpressor of salicylate hydroxylase (*35S-nahG*) (*105*), *sgt1b* (*106, 107*)) into *dcl4* to test the effects on autoimmune phenotypes. Most of these genetic backgrounds had no appreciable effect (Fig. S5A). The exception was mutation of the Hsp90 co-chaperone SGT1b which, similar to Hsp90 itself, has a dual role in regulation of R protein function. On the one hand, it is required for the function of some R proteins (*84, 108-110*), presumably by facilitating conformational changes required for oligomerization, and on the other hand, it is implicated in chaperone-assisted proteasomal degradation of many R proteins (*55, 56*). *sgt1b* enhanced the growth phenotype of *dcl4*, as it was stronger and reached 100% penetrance in *dcl4/sgt1b*, yet remained fully dependent on *DCL2* (Fig. 3B, Fig. S5B). Thus, in subsequent parts of our study, we used *dcl4/sgt1b* as a sensitized genetic background because its clearer phenotypic readout facilitated genetic dissection of DCL2-dependent autoimmunity.

## SUPPLEMENTAL TEXT 2

The strong, yet incomplete suppression of autoimmune phenotypes in *dcl4/sgt1b/l5* mutants (Fig. 3, Fig. S6) suggests that additional R proteins are involved in activation of immune responses. Clearly, RPP9 is not the only additional R protein, because *l5* and *rrp9* mutations did not show additive effects in suppression of *dcl4/sgt1b* (Fig. 3, Fig. S6). Furthermore, the appearance of novel autoimmune-like phenotypes in *dcl4/sgt1b/l5*, not seen in *dcl4/sgt1b* (Fig. S6), suggests that mutation of *L5* itself may condition further R protein activation. We consider two broad, and not mutually exclusive scenarios as likely explanations for this observation. First, since *L5* is also the host gene for the microRNA miR472 that targets several CNL-encoding mRNAs (*111*), it is possible that increased CNL expression in *l5-miR472* mutants occasionally triggers autoimmunity. Second, since L5 is conserved across *Arabidopsis thaliana* ecotypes, similar to the well-known signaling node ATR1-L1/L2 (*76, 77*), it may be a target of pathogen effectors, and hence subjected to surveillance by other R proteins whose activation may therefore be promoted by genetic inactivation of *L5*.

