## Supplemental Figures for "Sensing of viral RNA in plants via a DICER-LIKE Ribonuclease"

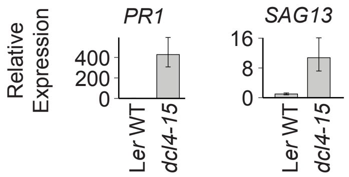


**Supplemental Figure 1. Defense response genes are upregulated in L*er dcl4-15***
Quantitative RT-PCR analysis of relative accumulation of *PR1* and *SAG13* mRNA in 16-day-old seedlings of L*er* WT and *dcl4-15*. cDNA inputs were normalized to *ACTIN2* expression. y-axis shows fold change of expression relative to wild type.

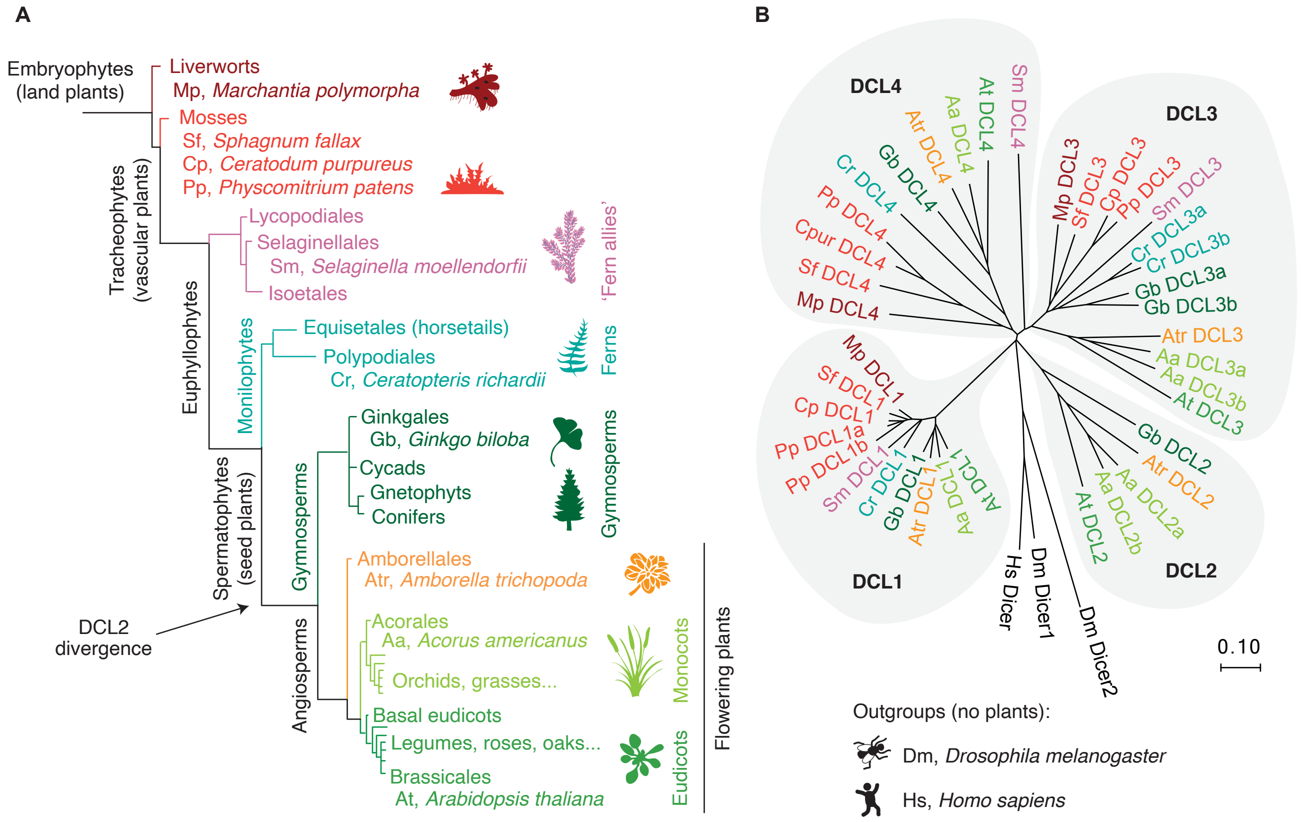


**Supplemental Figure 2. Phylogeny of plant DCL proteins**

(A) Phylogenetic relations of land plants highlighting the individuals from the major phylogenetic groups from which DCL protein sequences were retrieved.

(B) Phylogenetic tree of land plant DCL proteins based on multiple sequence alignment produced by Clustal-Ω (see Methods). The length of the branches represents the evolutionary distance in numbers of amino acid substitutions per site. While DCL1, DCL3 and DCL4 are found in all land plants, DCL2 is only found in spermatophytes.

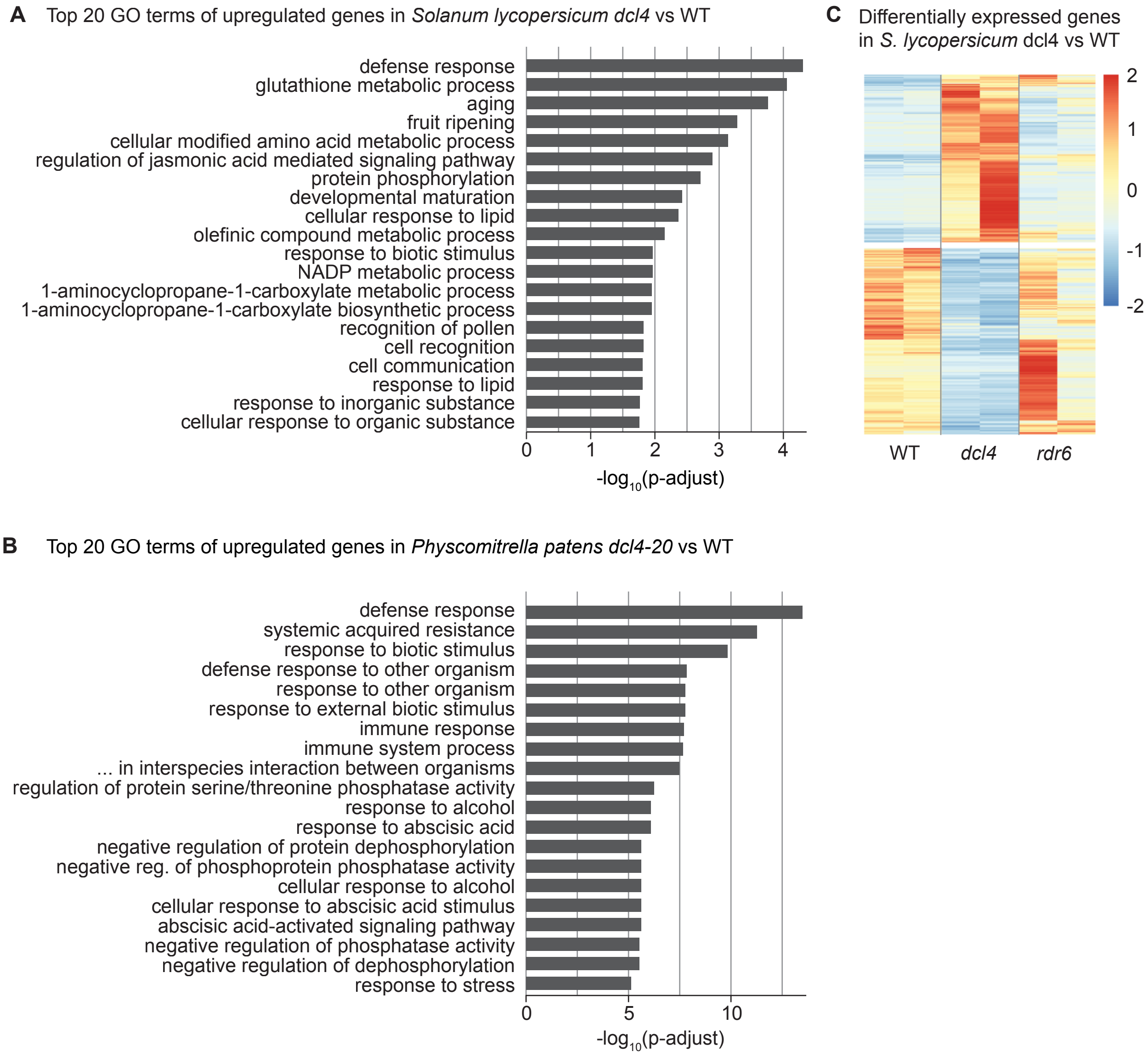


**Supplemental Figure 3. Enrichment of functionally related genes in sets of genes upregulated in *Solanum lycopersicum* and *Physcomitrium patens dcl4* mutants**

(A) Top 20 gene ontology (GO) terms enriched in genes significantly upregulated in *S. lycopersicum dcl4* compared to wild type, as determined by mRNA-seq.

(B) Top 20 gene ontology (GO) terms enriched in genes significantly upregulated in *P. patens dcl4-20* compared to wild type, as determined by mRNA-seq.

(C) Row-normalized heat map showing expression in *S. lycopersicum* wild type, *dcl4* and *rdr6*, of genes differentially expressed between *dcl4* and wild type.

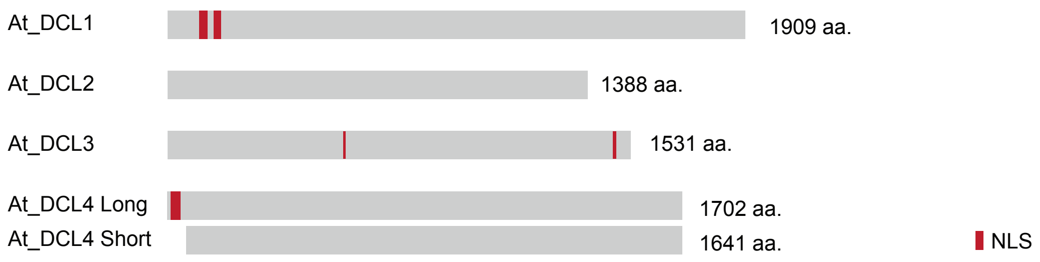


**Supplemental Figure 4. Presence of predicted nuclear localization signals in Arabidopsis DCL proteins**
Schematic representation of Arabidopsis DCL1-4 proteins. The positions of predicted nuclear localization signals (NLSs), as determined with the NLS prediction software NLS mapper, are highlighted in red. Please notice that *DCL4* is transcribed from two alternative transcription start sites of which only the minor, long form contains an NLS.

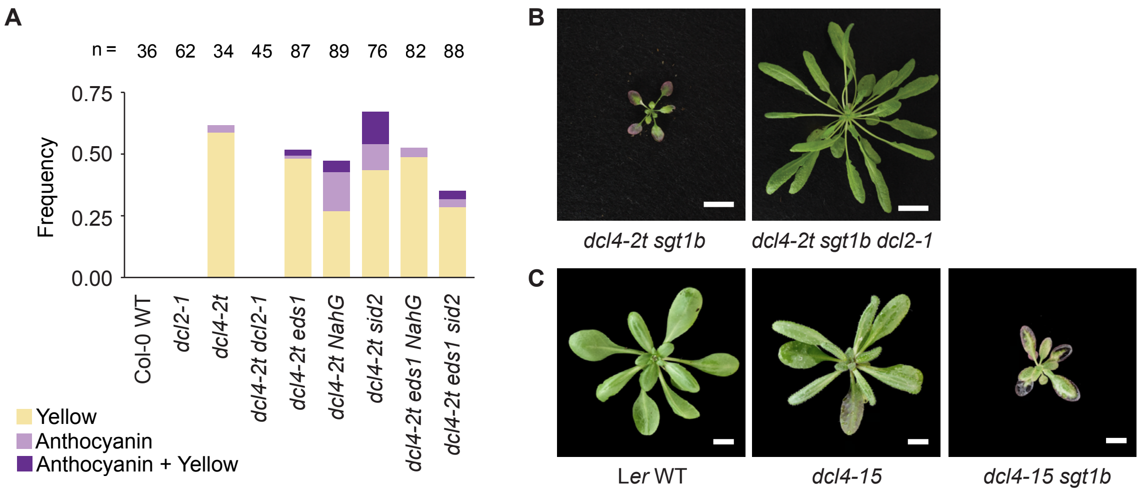


**Supplemental Figure 5. Visible effects of immune signaling and *sgt1b* mutants on leaf phenotypes of *dcl4* mutants**
(A) Frequency of plants showing either leaf yellowing (yellow), visible anthocyanin accumulation (anthocyanin), or both (yellow + anthocyanin) in the indicated genotypes (all in the Col-0 ecotype). Plants were grown in long days, first on sterile MS plates, and healthy seedlings at day 11 were transferred to soil and grown for 17 additional days.
(B) Rosette phenotypes of *dcl4-2t/sgt1b* and *dcl4-2t/sgt1b/dcl2-1* assesed after 49 days of growth in short days from soil-germinated seeds. Size bar, 2 cm.
(C) Representative photographs of 3-week old rosettes of the indicated genotypes in the L*er* ecotype. As in accession Col-0, the penetrance of the rosette phenotype of *dcl4-15/sgt1b-1* was 100%.

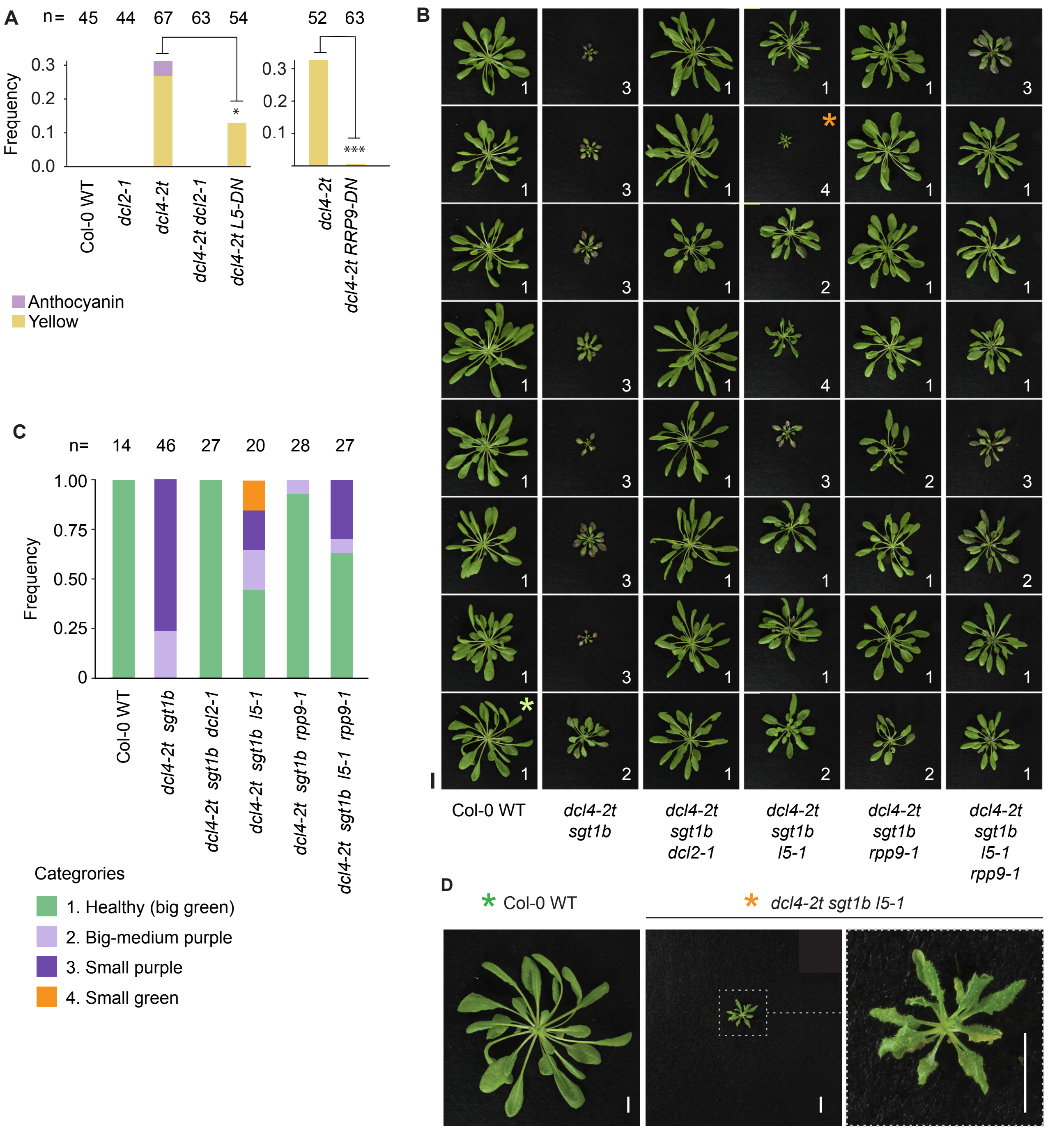


**Supplemental Figure 6. Genetic interactions between *DCL4*, *L5* and *RPP9***

(A) Quantification of rosette phenotypes indicating the fraction of plants showing visible leaf yellowing (yellow) or anthocyanin accumulation (anthocyanin). DN, dominant negative. The experiments shown use one line (out of many generated) of *dcl4-2t/L5-DN* (left) and of *dcl4-2t/RPP9-DN* (right) propagated to homozygosity for the transgene encoding the dominant negative R protein. Asterisks indicate significance of difference to *dcl4-2t* ($\chi^2$ test), * $P < 0.05$, *** $P < 0.001$.

(B) Representative individuals exemplifying the degree and variability of suppression of the *dcl4* growth phenotype by additional mutation in the *R* genes *L5* and *RPP9* and their combination, compared to that of *DCL2* mutation. Numbers on the lower-right corner of each photograph indicate the category assigned to each plant as described in C. The two photographs marked with asterisks on the right-upper corner are magnified in D. Scale bar, 2 cm.

(C) Quantification of rosette phenotypes of the plants photographed in B and additional individuals of each genotype (see n numbers), according to categories 1-4 as described.

(D) Magnification of the photographs marked with asterisks in panel B, to highlight the autoimmune-like phenotype with severely stunted growth but no visible leaf chlorosis exhibited by a small subset (< 5%) of individuals in *dcl4-2t/sgt1b/l5* populations (category 4 in B and C). Such phenotypes were not observed in *dcl4-2/sgt1b* populations in either accession Col-0 or L*er*. Size bars, 1 cm.
